# Genome-wide methylation profiling of cell-free DNA in maternal plasma using Methylated DNA Sequencing (MeD-seq)

**DOI:** 10.1101/2024.08.29.610227

**Authors:** Marjolein M. van Vliet, Ruben G. Boers, Joachim B. Boers, Olivier J.M. Schäffers, Lotte E. van der Meeren, Régine P.M. Steegers-Theunissen, Joost Gribnau, Sam Schoenmakers

**Affiliations:** Department of Obstetrics and Gynaecology, Erasmus MC, the Netherlands; Department of Developmental Biology, Erasmus MC, the Netherlands; Department of Pathology, Erasmus Medical Centre Rotterdam, the Netherlands; Department of Pathology, Leiden University Medical Center, the Netherlands

## Abstract

**Background:** Placental-originated cell-free DNA (cfDNA) provides unique opportunities to study (epi)genetic placental programming remotely, but studies investigating the cfDNA methylome are scarce and usually technologically challenging. Methylated DNA sequencing (MeD-seq) is well-compatible with low cfDNA concentrations and has a high genome-wide coverage. We therefore aim to investigate the feasibility of genome-wide methylation profiling of first trimester maternal cfDNA using MeD-seq, by identifying placental-specific methylation marks in cfDNA.

**Methods:** We collected cfDNA from non-pregnant controls (female n=6, male n=12) and pregnant women (n=10), first trimester placentas (n=10), and paired preconceptional and first trimester buffy coats (total n=20). Differentially methylated regions (DMRs) were identified between pregnant and non-pregnant women. We investigated placental-specific markers in maternal cfDNA, including *RASSF1* promoter and Y-chromosomal methylation, and studied overlap with placental and buffy coat DNA methylation.

**Results:** We identified 436 DMRs between cfDNA from pregnant and non-pregnant women which were validated using male cfDNA. *RASSF1* promoter methylation was higher in maternal cfDNA (fold change 2.87, unpaired t-test p<0.0001). Differential methylation of Y-chromosomal sequences could determine fetal sex. DMRs in maternal cfDNA showed large overlap with DNA methylation of these regions in placentas and buffy coats, indicating a placental and immune-cell contribution to the pregnancy-specific cfDNA methylation signature. Sixteen DMRs in maternal cfDNA were specifically found only in placentas. These novel potential placental-specific DMRs were more prominent than *RASSF1*.

**Conclusions:** MeD-seq can detect (novel) genome-wide placental DNA methylation marks and determine fetal sex in maternal cfDNA. This study supports future research into maternal cfDNA methylation using MeD-seq.

**Graphical abstract:** 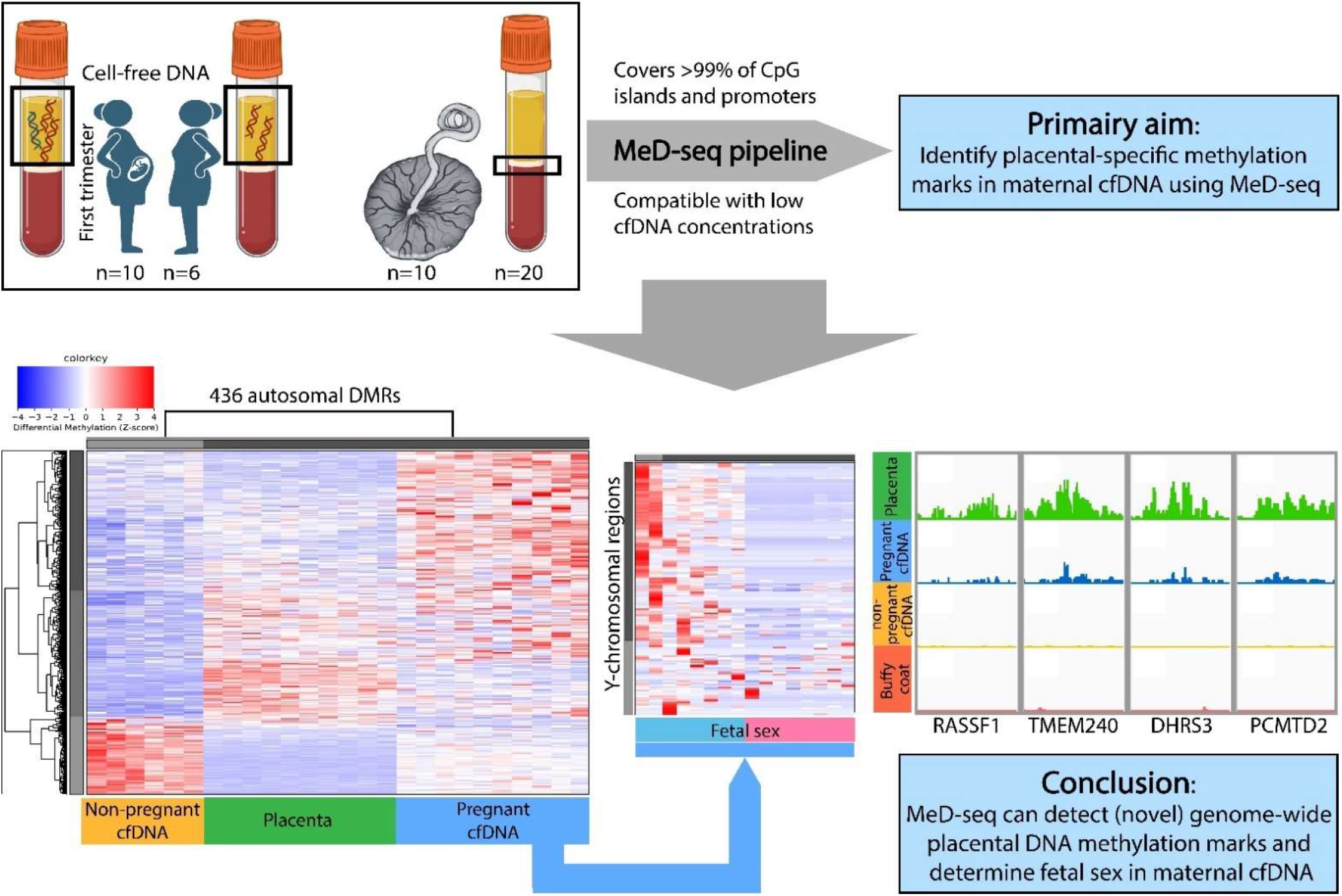

Studies investigating the maternal cell-free DNA (cfDNA) methylome are scarce and generally technologically challenging. We identified 436 autosomal differentially methylated regions (DMRs) between cfDNA from pregnant and non-pregnant women, using the innovative methylated DNA sequencing (MeD-seq) technique. Y-chromosomal methylation could determine fetal sex, we show hypermethylation of the placental-marker *RASSF1*, and identify 16 novel placental-specific markers in maternal cfDNA including DMRs related to *TMEM240, DHRS3*, and *PCMTD2*. This pilot study supports future research into the maternal cfDNA methylome using MeD-seq.

## Background

A well-functioning placenta is essential for healthy fetal development, but the placenta is largely inaccessible during gestation. The direct study of placental tissue necessitates invasive proceduresWilson, Gagnon (1), or postpartum placental examination. Therefore, less invasive techniques to study placental development throughout gestation are warranted.

Turnover and shedding of placental cells result in release of placental-originated cell-free DNA (cfDNA) in the maternal blood stream. Maternal plasma-derived cfDNA is widely used to perform non-invasive prenatal testing (NIPT), enabling screening for chromosomal abnormalities (2). However, delving into the epigenetic landscape of maternal cfDNA could allow for applications beyond current practice (3). DNA methylation is an important epigenetic mechanism playing a crucial role in placental growth, development, and function by regulating gene-expression (4-6). Differences in placental DNA methylation have been associated with gestational age and adverse obstetrical outcomes including preeclampsia (7-9). Maternal plasma-derived cfDNA could facilitate the non-invasive study of placental DNA methylation in health and disease already from the first trimester onwards.

A major challenge in using maternal cfDNA as a proxy for placental DNA is the relatively low contribution of placental-derived cfDNA to total maternal cfDNA. The largest contributor to maternal cfDNA is the maternal hematopoietic system, with a placental-derived fraction of roughly 10% at the end of first trimester (10-12). However, since DNA methylation is highly cell type specific, cfDNA methylation profiles can aid in determining their tissue of origin (4, 13). Previous studies identifying placental-specific DNA methylation markers mostly focused on methylation differences on chromosomes 13, 18, and 21, which have been shown to be potentially useful in screening for chromosomal abnormalities (14-23), and/or focused on methylation differences between placental tissues and maternal whole blood (24-26). Nonetheless, our understanding of the genome-wide impact of pregnancy directly on the cfDNA methylome, and how this reflects changes in maternal and placental tissues is limited. The scarce studies on cfDNA are mostly performed with whole genome bisulfite sequencing (27-29), which is costly and technologically challenging due to degradation of already low DNA levels in cfDNA samples, or achieved a limited genome-wide coverage (30). At the same time, recent studies support the use of the cfDNA methylome as early predictive marker for preeclampsia development (31-33). This emphasizes the need to further explore the epigenetic landscape of cfDNA already during the early stages of pregnancy, ideally using less costly methods while ensuring a high genome-wide coverage.

We therefore used Methylated DNA sequencing (MeD-seq) to investigate its feasibility to study the maternal cfDNA methylome. MeD-seq is compatible with cfDNA and covers >50% of genome-wide cytosine-guanine dinucleotides (CpGs), detecting DNA methylation at >99% of all CpG islands and promoters (34, 35). We aim to investigate the impact of pregnancy on the first trimester maternal cfDNA methylome, and explore the placental and hematopoietic origin of identified differences. We further aim to identify placental-specific methylation marks in cfDNA. As proof-of-principle, we show the possibility to determine fetal sex in maternal cfDNA based on Y-chromosomal methylation marks.

## Methods

### Study design

We included pregnant women who participated in the Rotterdam Periconception Cohort (Predict study) at the Erasmus Medical Centre, Rotterdam. The Predict study is an ongoing prospective cohort study where women with a singleton pregnancy are recruited before 10 weeks of gestation and longitudinally followed throughout pregnancy (36, 37). Blood samples for cfDNA isolation were obtained around 11 weeks of gestation between November 2022 and March 2023.

Blood samples for cfDNA isolation from non-pregnant controls were previously obtained for research purposes from anonymous, healthy blood donors (HBDs) enrolled via the Dutch National blood bank (Sanquin). Additionally, we prospectively collected first trimester (around 9-12 weeks of gestation) placental tissues after elective surgical abortions at a Dutch abortion clinic between July and August 2023. All pregnancies had to be without known congenital abnormalities. Lastly, maternal paired buffy coats were collected from women who participated within the Predict study both before pregnancy (preconception) as in the first trimester of a subsequent pregnancy.

Both the Predict study (MEC-2004-227) and the research protocol to collect placental tissues (MEC-2022-0788) were approved by the Medical Ethics Committee of the Erasmus Medical Centre. All participants provided written informed consent.

### Processing of samples

Blood samples for cfDNA isolation were collected in CellSave preservative tubes. Plasma was subsequently isolated by two centrifugation steps (10 minutes at 1,711g — 2,000g followed by 10 minutes 12,000g — 16,000g) (38). Blood samples for DNA isolation of buffy coats were collected in EDTA tubes. Buffy coats were isolated after centrifugation of the blood sample (10 minutes at 2,000g). Plasma and buffy coats were stored at −80 °C prior to (cf)DNA isolation. Placental tissues were stored in buffered 4% formaldehyde solution.

### (cf)DNA isolation

cfDNA was isolated from 4 ml of plasma using the QIAamp Circulating Nucleic Acid Kit. DNA extraction and isolation from buffy coats were performed using the TECAN freedom EVO robot combined with the Promega ReliaPrep™ Large Volume HT gDNA Isolation System. Placental samples were formalin-fixed and full thickness biopsies were taken followed by embedding in formalin-fixed paraffin-embedded (FFPE) blocks. Four to six slides of 4-6 μm were cut. One slide per sample was stained using hematoxylin and eosin (H&E) and placental parenchyma was evaluated by a perinatal pathologist (LEM). DNA was extracted and isolated from subsequent slides with normal first trimester placental parenchyma using the QIAamp DSP DNA FFPE Tissue kit according to the manufacturer’s protocol.

### MeD-seq assay

MeD-seq assays were performed as previously described (34, 35, 39). Genomic DNA and plasma-derived cfDNA were digested with the methylation-dependent restriction enzyme LpnPI (New England Biolabs, Ipswich, MA) generating 32 bp DNA fragments containing the methylated CpG in the middle. Samples were sequenced on the Illumina NextSeq2000 platform.

### Data analysis

Differentially methylated regions (DMRs) were detected using a previously established bioinformatics pipeline (34, 35, 39). Custom python scripts were used to process acquired DNA methylation profiles. Raw fastq files were subjected to Illumina adaptor trimming and reads were filtered based on LpnPI restriction site occurrence between 13–17 bp from either the 5′ or 3′ end of the read. Reads that passed the filter were mapped to hg38 using bowtie2 and BAM files were generated using SAMtools version 0.1.19. Genome-wide individual LpnPI site scores were used to generate read count scores for: transcription start sites (TSS, 1 kb before and 1 kb after), CpG-islands, and gene bodies (1 kb after TSS till TES). Gene and CpG-island annotations were downloaded from ENSEMBL (Homo_sapiens_hg38.GRCh38.79.gtf, www.ensembl.org).

Additionally, a sliding window technique was used to detect DMRs and the Chi-squared test on read counts was used for statistical testing. A Bonferroni corrected p-value ≤0.05 was considered statistically significant. Z-score transformation of the read count data was applied for unsupervised hierarchical clustering analyses.

DMRs were identified between cfDNA from pregnant and non-pregnant women. cfDNA from male HBDs were used as validation. Y-chromosomal methylated DNA sequences were identified between cfDNA from male and female HBDs and subsequently applied to maternal cfDNA to determine fetal sex. Methylation of the *RASSF1* promoter, known to be hypermethylated specifically in placental tissues (40), was compared between cfDNA from pregnant and non-pregnant women using an unpaired t-test standardized for number of reads per million (rpm). For DMRs identified in cfDNA, overlap with first trimester placental tissues and buffy coats was studied to explore tissue of origin of identified DMRs. To determine the presence or absence of cfDNA methylation signatures associated specifically with pregnant or with non-pregnant women, receiver operating characteristic (ROC) curves were calculated for each sample for each individual DMR. ROC curves of reference sets of pregnant women compared to non-pregnant women were used to calculate the optimal threshold (using the “scikit-learn” package Python) for each individual DMR. Samples above the threshold scored ‘1’, samples under the threshold scored ‘0’, leading to a binary score for each DMR. To investigate overlap of identified DMRs in cfDNA with placental tissues and with buffy coats, a cut-off value of ≥80% of samples scoring ‘1’ or ‘0’ was used for hypermethylated and hypomethylated DMRs, respectively.

## Results

An overview of used samples and their technical performance is provided in Table S1.

### DMRs between pregnant and non-pregnant cfDNA

To identify differences in cfDNA methylation related to pregnancy, we compared cfDNA from pregnant (n=10) and non-pregnant women (HBDs, n=6) (Figure 1A). For pregnant women, baseline characteristics are depicted in Table 1, and cfDNA was collected between 10+5 – 11+5 weeks of gestation (Figure 1B). Using MeD-seq (Figure 1C), we identified 436 autosomal DMRs with a fold change (FC) ≥2, of which 338 (77.5%) were hypermethylated in pregnant women (Table S2). Unsupervised hierarchical clustering using identified DMRs shows clear clusters between pregnant and non-pregnant women (Figure 1D). As a validation, we show large overlap of male cfDNA (n=12) with non-pregnant women showing robustness of our results (Figure S1).

**Table 1.** Baseline characteristics of pregnant women with cfDNA samples (n=10). Categorical variables are described using absolute numbers and percentages (%). Continuous variables are described using medians and interquartile ranges (IQR).IVF/ICSI = in vitro fertilization/intracytoplasmic sperm injection.

|  |  |
| --- | --- |
| **Maternal age (years)** | 32.4 (30.7 – 34.3) |
| **BMI at study entry** | 25.1 (22.8 – 26.8) |
| **Country of birth** |  |
| Dutch | 8 (80%) |
| Non-Dutch | 2 (20%) |
| **Nulliparous** | 6 (60%) |
| **Smoking during pregnancy** | 1 (10%) |
| **Mode of conception** |  |
| Spontaneous | 4 (40%) |
| IVF/ICSI | 6 (60%) |
| **Folic acid supplement use** | 10 (100%) |
| Started preconception | 9 (90%) |
| **Male fetus** | 5 (50%) |

**Figure 1.**
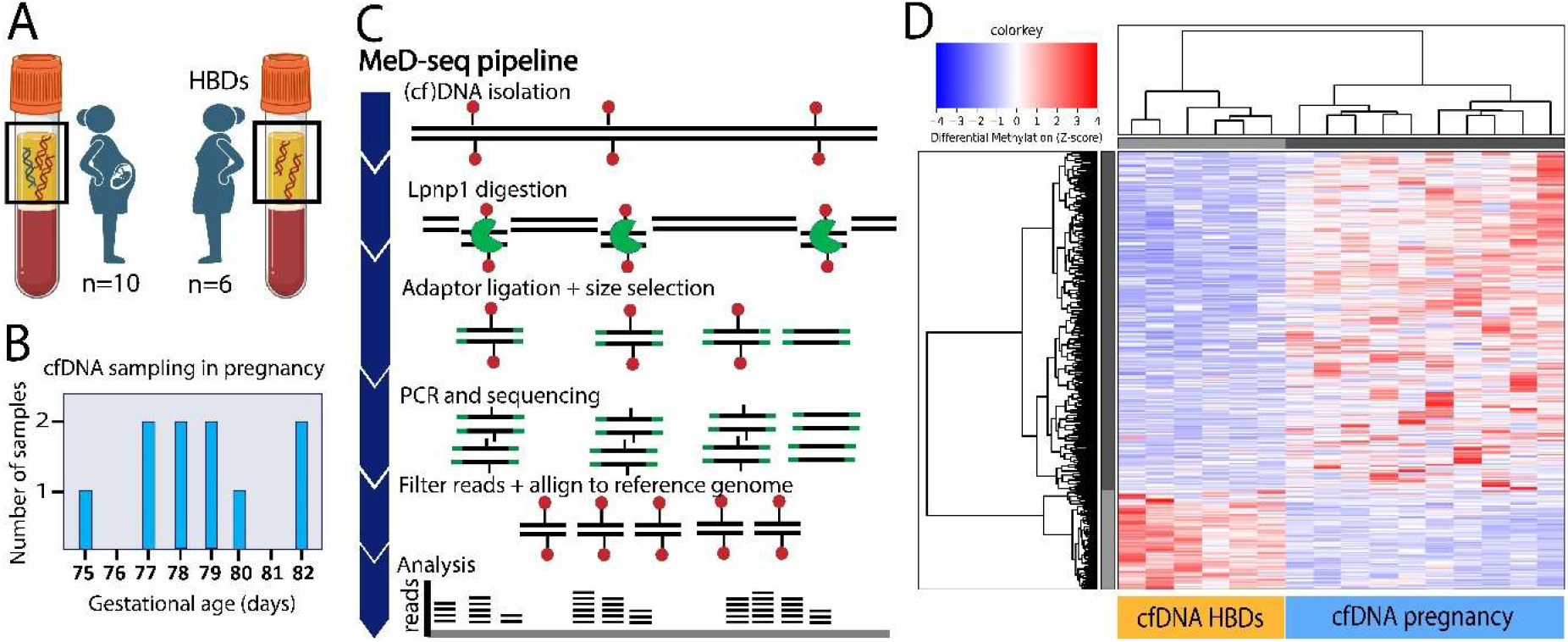
Identification of DMRs between cfDNA from pregnant (first trimester) and non-pregnant women (HBDs). A)Study population consists of pregnant women at the end of the first trimester and non-pregnant women. cfDNA in plasma from pregnant women partly originates from the placenta. B)Gestational age at time of cfDNA sampling in pregnancy. C)Schematic overview of the MeD-seq pipeline. (cf)DNA is digested by LpnPI which recognizes methylated CpGs and cuts the DNA 16 bp up-and downstream leading to 32 bp fragments. This is followed by adaptor ligation, size selection of cfDNA fragments (32 bp), amplification and sequencing using the Illumina NextSeq2000 platform. Reads are filtered based on a central methylated CpG and aligned to a reference genome (hg38) prior to analysis. D)Heatmap showing an unsupervised hierarchical clustering using autosomal DMRs with a Fold Change ≥2 between cfDNA from pregnant women in first trimester and non-pregnant women (HBDs) showing clear separation between the two groups. Z-scores on normalized MeD-seq data are used to visualize the data, red represents hypermethylation, blue represents hypomethylation.

### Placental origin of methylation profiles in cfDNA from pregnant women

To explore which DMRs in maternal cfDNA could be of placental origin, we compared DMRs identified in cfDNA with MeD-seq data from first trimester placental tissues (n=10) (Figure 2A). Placental tissues were collected between 9+2 – 12+2 weeks of gestation (Fig 2B, Table S1). Unsupervised hierarchical clustering analysis shows that first trimester placental tissues cluster between cfDNA from pregnant and non-pregnant women (Figure 2C). Based on our cumulative methylation score, the majority of DMRs hypermethylated in cfDNA from pregnant women were in general also detected in placental tissues, while DMRs hypermethylated in cfDNA from non-pregnant women were not found in placental tissues (Figure 2D, Table S3). More strictly, of 338 DMRs hypermethylated and 98 DMRs hypomethylated in cfDNA from pregnant women, 143 (42.3%) and 88 (89.8%) were respectively consistently also hypermethylated or hypomethylated in ≥80% of placental tissues (Table S3). DMRs that are hyper-or hypomethylated in both maternal cfDNA and placental tissues, as compared to cfDNA from non-pregnant women could indicate a placental-origin of these DMRs in maternal cfDNA. A well-studied placental-specific DNA methylation mark is hypermethylation of the *RASSF1* promoter (41). As expected, the *RASSF1* promoter displayed a higher methylation level in first trimester placental tissues as compared to cfDNA from pregnant or non-pregnant women (Figure 2E-F, Table S4). Moreover, the *RASSF1* promoter displayed a higher methylation level in cfDNA from pregnant as compared to non-pregnant women (FC 2.87, unpaired t-test p<0.0001) (Figure 2E-F, Table S4), although this did not remain statistically significant in our genome-wide analyses after Bonferroni correction.

**Figure 2.**
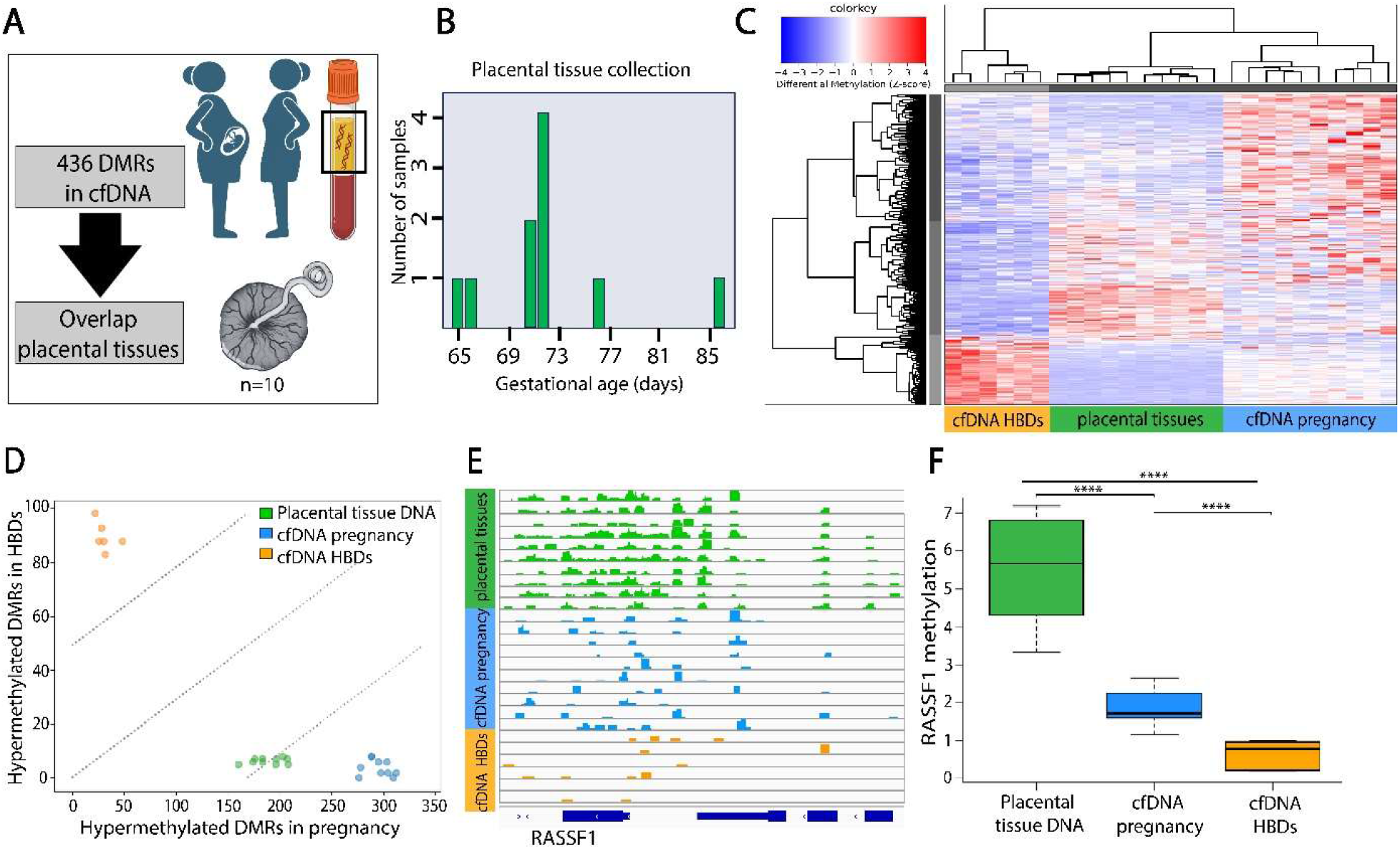
Placental origin of methylation profiles in cfDNA from pregnant women. A)DMRs were identified in cfDNA between non-pregnant women (HBDs) and pregnant women in the first trimester. For identified DMRs, overlap with first trimester placental tissue biopsies is studied. B)Gestational age at time of placental tissue collection. C)Heatmap visualizing unsupervised hierarchical clustering of first trimester placental tissue biopsies between cfDNA from pregnant women and non-pregnant women (HBDs). Selected autosomal DMRs were identified between pregnant and non-pregnant women (HBDs). Red represents hypermethylation, blue represents hypomethylation. D)After generating a cumulative methylation score, most DMRs hypermethylated in cfDNA from pregnant women are also hypermethylated in first trimester placental tissue biopsies (x-axis). DMRs specifically hypermethylated in cfDNA from non-pregnant women (HBDs) are not hypermethylated in first trimester placental tissue biopsies (y-axis). Each dot represents one sample. E,F)Gene-tracks and boxplots show hypermethylation of the *RASSF1* promoter in placental tissue biopsies, and higher methylation of the *RASSF1* promoter in cfDNA from pregnant women compared to cfDNA from non-pregnant women (HBDs) (t-test p<0.0001). *In our genome-wide analysis, the difference in *RASSF1* promoter methylation between cfDNA from pregnant and non-pregnant women did not remain statistically different after Bonferroni correction.

### Fetal sex determination based on DNA methylation

To assess the feasibility of determining fetal sex using cfDNA methylation profiles, we first identified 147 Y-chromosomal regions containing DNA methylation with a FC ≥2 in cfDNA from male (n=6) compared to female HBDs (n=6) (Figure 3A, Table S5). Using identified Y-chromosomal regions, two clear clusters were identified in maternal cfDNA (Figure 3B). Women bearing boys (n=5) showed hypermethylation at numerous Y-chromosomal regions as compared to women pregnant with girls (n=5) (Figure 3B-C). Epigenetic marks can thus be used to determine fetal sex in maternal cfDNA. The minimal detection of Y-chromosomal reads in women pregnant with girls (Figure 3B-C), can be explained by mapping errors caused by large regions displaying high levels of homology between parts of the X-and Y-chromosome.

**Figure 3.**
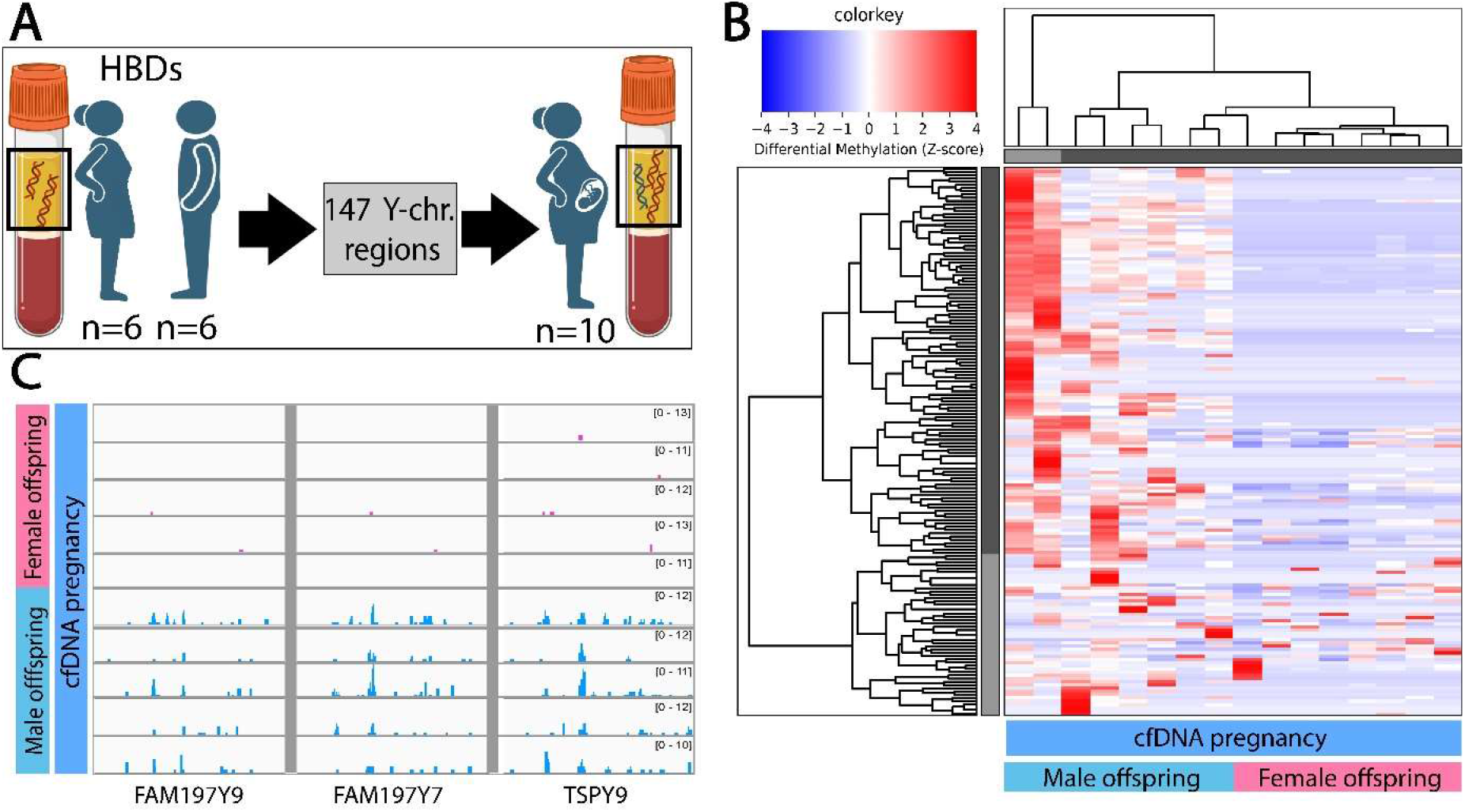
Fetal sex determination in cfDNA from pregnant women. A)Y-chromosomal regions containing DNA methylation were identified between male and female healthy blood donors (HBDs). Identified Y-chromosomal regions were studied in cfDNA from pregnant women. B)Heatmap visualizing unsupervised hierarchical clustering of cfDNA from pregnant women based on identified Y-chromosomal regions shows clear clusters of women bearing male offspring (blue bar, left) and female offspring (pink bar, right). Red represents hypermethylation, blue represents hypomethylation. C)Example gene-tracks of Y-chromosomal regions in cfDNA from pregnant women showing methylation only in women bearing male offspring.

### Overlap identified DMRs with buffy coats and novel placental-specific markers

Not all DMRs identified in maternal cfDNA were detected in placental tissues (Figure 2A-B, Table S3). This could indicate pregnancy-induced changes in maternal cfDNA from other tissues. During pregnancy, physiological changes result in an increase in white blood cells while simultaneously a change in composition of maternal immune cells occurs (43). Since most cfDNA originates from maternal hematopoietic cells (10), we hypothesized that pregnancy-induced physiological immunological changes might explain part of the observed methylation differences between cfDNA from pregnant and non-pregnant women.

To first identify DNA methylation changes in buffy coats related to pregnancy, we compared DNA methylation profiles between paired buffy coats collected preconception (n=10) and in the first trimester of a subsequent pregnancy (n=10) (Figure 4A). Baseline characteristics of included women are depicted in Table S6. In contrast to cfDNA, we identified no DMRs between preconception and first trimester buffy coats.

**Figure 4.**
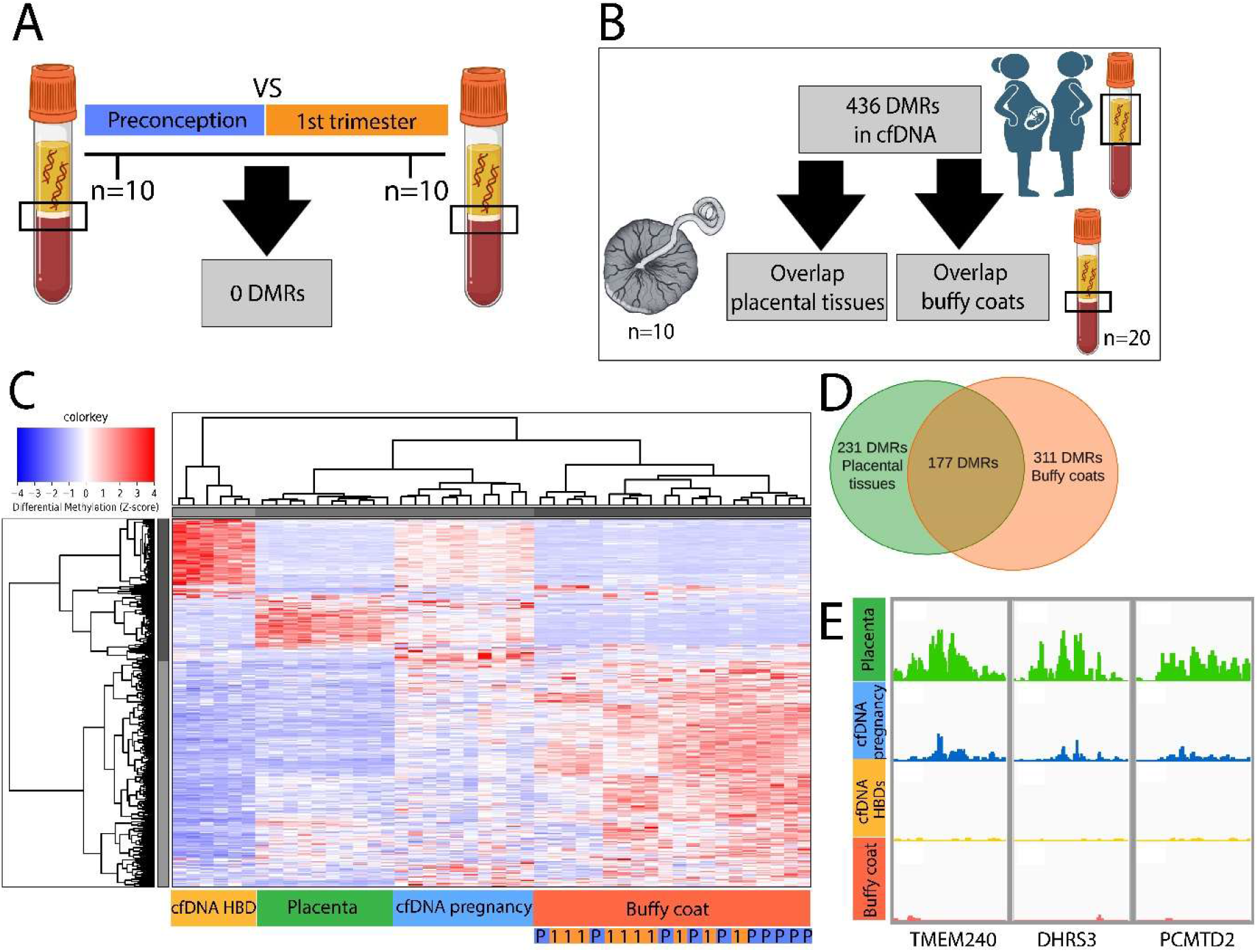
Origin of DMRs identified in cfDNA: overlap with placental tissues and buffy coats and identification of novel placental-markers in cfDNA. A)No DMRs were found between paired buffy coats collected preconception and in the first trimester of pregnancy. B)Overlap with DNA methylation in placental tissue biopsies and pooled buffy coats was studied for DMRs identified in cfDNA. C)Heatmap visualizing unsupervised hierarchical clustering for DMRs found between cfDNA from pregnant women and non-pregnant women (HBDs), shows larger overlap of buffy coats with cfDNA from pregnant women as with cfDNA from non-pregnant women. Selected autosomal DMRs were identified with a Fold Change ≥2. Red represents hypermethylation, blue represents hypomethylation. Buffy coats collected during the first trimester pregnancy (P, blue) cluster between buffy coats collected preconceptionally (1, orange). D)Venn diagram visualizing overlap of DMRs identified in cfDNA from pregnant women compared to non-pregnant women with first trimester placental tissue biopsies and buffy coats. Used cutoff is DMRs present in ≥80% of placental tissue biopsies or buffy coats based on our cumulative methylation score (Table S3). E)Example gene-tracks of potential placental-specific DMRs in maternal cfDNA. Gene-tracks for DMRs with highest fold change between cfDNA from pregnant and non-pregnant women. DMRs are hypermethylated in cfDNA from pregnant women and placental tissue biopsies as compared to cfDNA from non-pregnant women (HBDs). DMRs corresponding to *TMEM240, DHRS3*, and a CpG island upstream of *PCMTD2* show no hypermethylation in buffy coats indicative of a placental-origin.

To further explore possible origins of DMRs identified in cfDNA, next to overlap with placental tissues, we studied overlap of identified DMRs with pooled buffy coats (n=20) (Figure 4B). Surprisingly, the majority of DMRs identified in maternal cfDNA were also present in buffy coats (Figure 4C-D). Based on our cumulative methylation score, 226 (66.9%) of DMRs hypermethylated and 85 (86.7%) of DMRs hypomethylated in maternal cfDNA were respectively also hyper-or hypomethylated in ≥80% of buffy coat samples (Table S3).

While 231 (53.0%) of DMRs identified in maternal cfDNA were present in ≥80% of placental tissue biopsies, 177 of these were also present in ≥80% of buffy coat samples (Figure 4D, Table S3). These DMRs are therefore not placental-specific, but could also be of hematopoietic origin. The large overlap of DMRs in maternal cfDNA with DNA methylation of buffy coats might reflect an overall increased immune response during pregnancy.

DMRs in maternal cfDNA that overlap with DNA methylation in placental tissues but not with buffy coats, are potential placental-specific DMRs. We identified 16 DMRs that were hypermethylated in ≥80% of both maternal cfDNA and placental tissue samples, while hypomethylated in both ≥80% of HBD cfDNA and buffy coat samples (Table S3). These DMRs are therefore likely to represent placental-specific markers in cfDNA. To illustrate, Figure 4E shows gene tracks for 3 DMRs with the highest FC between cfDNA from pregnant and non-pregnant women. The three top-ranked DMRs correspond to *TMEM240* (FC 10.2), *DHRS3* (FC 7.6), and a CpG island 40kb upstream of *PCMTD2* (FC 6.5).

## Discussion

We applied MeD-seq to investigate the genome-wide impact of first trimester pregnancy on the maternal plasma-derived cfDNA methylome. We identified numerous DMRs between cfDNA from pregnant and non-pregnant women. Next, we confirmed identification of placental-specific methylation marks in maternal cfDNA by showing hypermethylation of the *RASSF1* promoter – a known placental-specific marker – and the possibility to determine fetal sex based on Y-chromosomal DNA methylation. Moreover, we identified novel potential placental-specific methylation marks in maternal cfDNA, such as hypermethylated DMRs in *TMEM240, DHRS3*, and a CpG island upstream of *PCMTD2*, that were more robust as compared to *RASSF1* hypermethylation.

Although the majority of DMRs in maternal cfDNA overlapped with placental DNA methylation, most DMRs also overlapped with DNA methylation of buffy coats. This suggests a more prominent immune-cell type(s) related contribution to DMRs in cfDNA from pregnant as compared to non-pregnant women. This is in line with other studies that recently found that the increased cfDNA levels in patients with several types of cancers is predominantly caused by increased levels of leukocyte-derived cfDNA (42, 43), as is also found for patients with acute pancreatitis or sepsis. Our findings therefore suggest that the cfDNA methylation profile reflects an overall increased immune response during pregnancy (10, 44).

DMRs in maternal cfDNA that were not shared with either placental tissues or buffy coats, could be attributable to physiological pregnancy-related changes in other (high-turnover) maternal tissues that contribute to cfDNA, such as liver-or endothelium (10). Another explanation could be a different contribution of hematopoietic and placental cell types to buffy coats or tissue biopsies as compared to their contribution to cfDNA. For example, in placental tissue biopsies, DNA methylation reflects the average methylation of all cells in the selected sample, while for cfDNA the contribution of different placental cell types could be influenced by apoptosis rate and anatomical position in relation to the maternal blood stream.

Although fetal sex can already be determined in maternal cfDNA based on the presence of Y-chromosomal genetic material (2), we were able to establish the fetal sex by combining numerous methylated Y-chromosomal regions in maternal cfDNA. These findings suggest the potential use for specific DMRs on other chromosomes to aid in the detection of known genetic disorders as previously shown for aneuploidies, (15, 16, 21-23) and also for chromatinopathies affecting genome-wide DNA methylation (45). Combining genetic and epigenetic information may further improve the current practice of NIPT and the potential added value of combining both sources warrants further study.

MeD-seq can be used to discover genome-wide DMRs directly in cfDNA, instead of using differences between whole blood and placental tissues. This may more reflect the turnover of specific cell types and can lead to additional markers relevant for cfDNA applications in pregnancy. For example, the previously mentioned hypermethylation of DMRs in *TMEM240, DHRS3*, and a CpG island upstream of *PCMTD2* were more robust in cfDNA of pregnant compared to non-pregnant women, as compared to *RASSF1* hypermethylation. Application of the UCSC database shows enrichment for regulatory regions (ReMap database) for two of our DMRs (*DHRS3* and *PCMTD2*), possibly related to a functional role for DNA methylation in gene regulation of nearby genes. The three top-ranked DMRs show enrichment for binding of numerous proteins, mainly including BRD4 which is involved in regulating cell differentiation, and ESR1 and CTCF, which play a role in transcription regulation. Additionally, comparable to hypermethylation of *RASSF1*, hypermethylation of *TMEM240* and *DHRS3* have been described in several types of cancers and these genes have been proposed as tumor suppressor genes (46-48). Similarities in DNA methylation between the placenta and cancers are well known, and are suggested to be involved in their mutual properties such as proliferation, cell invasion, and immune modulation (49).

A previous study identified differentially methylated CpGs between cfDNA from pregnant and non-pregnant women (30). They found 8277 differently methylated CpGs, of which 1704 CpGs, corresponding to 501 genes, were hypermethylated in cfDNA from pregnant compared to non-pregnant women, as well as in chorionic villus samples compared to maternal leukocytes. Although we focused on DMRs instead of individual CpGs, 11 of these identified genes overlapped with genes corresponding to our identified DMRs, of which for 6 the previously identified CpGs were captured by our DMRs: *DCAF10, DENND2D, OSR2, RNF126P1, TMEM17*, and *TMEM233* (Figure S2). These DMRs are therefore also likely robust placental-specific DNA methylation markers in maternal cfDNA.

In contrast to this previous study, most DMRs identified in our study were hypermethylated instead of hypomethylated in cfDNA from pregnant as compared to non-pregnant women. This is likely due to technological differences. We selected DMRs with a fold change ≥2 to reduce the risk of false positive DMRs. Consequently, MeD-seq is biased towards hypermethylated regions. For example, for regions highly methylated in blood cells and thus in cfDNA, but hypomethylated in placental tissues, the relative decrease in methylation of maternal cfDNA caused by hypomethylated placental-originated cfDNA fragments will often be below 2 and remain undetected.

The relatively low contribution of placental-derived cfDNA remains a major challenge when using cfDNA as a proxy for placental DNA (11, 12). In future studies, fragment size of cfDNA may be used to enrich for placental-originated cfDNA, since placental-originated cfDNA fragments are shorter than maternal cfDNA fragments (50). Combining DNA methylation analyses with other cfDNA features, referred to as “fragmentomics”, could improve tissue of origin identification and analyses (50, 51). Importantly, the contribution of different placental cell types (e.g. syncytiotrophoblast, cytotrophoblast, extravillous trophoblast) to cfDNA is currently unclear and probably also changes during gestation, driven by developmental changes in the placenta composition itself, as is observed for cell-free RNA (52). Therefore, future studies should strive to establish DNA methylation profiles for specific placental cell type as reference.

Recent studies have shown promising results for the use of the cfDNA methylome as predictor of obstetric complications, such as preeclampsia (32, 33). More research is warranted to establish robust, easy to use biomarkers for obstetric health based on the cfDNA methylome. Ideally, early prenatal genetic screening and application of identified biomarkers can in the future jointly be incorporated in a single test. The combined study of (epi)genetics in maternal cfDNA could expand the current NIPT to what we would call an Epigenome-wide Non-Invasive prenatal Test (Epi-NIPT).

This study is one of the first to study genome-wide methylation of cfDNA in pregnant women. As compared to other techniques, MeD-seq could be a less costly and easier to perform method for low concentration cfDNA methylation profiling while ensuring a high genome-wide coverage. However, limitations of this pilot study are that we did not correct for potential confounders such as age, ethnicity, and smoking which can impact DNA methylation (53-55). For HBDs and placental tissues, we had no access to detailed participant’s characteristics. Furthermore, MeD-seq has a bias towards hypermethylated regions and the highly fragmented nature of cfDNA may influence the coverage of the genome. Lastly, differences in placental cell type contributions in tissue biopsies and cfDNA could have distorted our results. Future studies should therefore aim to establish reference methylation profiles for different placental cell types.

## Conclusion

Research into the cfDNA methylome could allow for cfDNA applications beyond current practice, however, exploring the epigenetic landscape of cfDNA during pregnancy is still in its infancy. This pilot study revealed feasibility of genome-wide methylation profiling of maternal cfDNA using the MeD-seq technology. As a proof-of-concept, we showed that genome-wide placental DNA methylation marks can be identified in first trimester maternal cfDNA and fetal sex can be determination based on Y-chromosomal DNA methylation. The added value of epigenome-wide analyses of cfDNA using MeD-seq in the current practice of prenatal genetic testing and in relation to obstetric complications and gestational age warrants further study.

## Availability of data and materials

The MeD-seq datasets supporting the conclusions of this article are available in the Sequence Read Archive (SRA) repository at the National Center for Biotechnology Information, with accession number PRJNA1108949 via:https://www.ncbi.nlm.nih.gov/sra/?term=PRJNA1108949.

---

**Supplemental Figure1.**
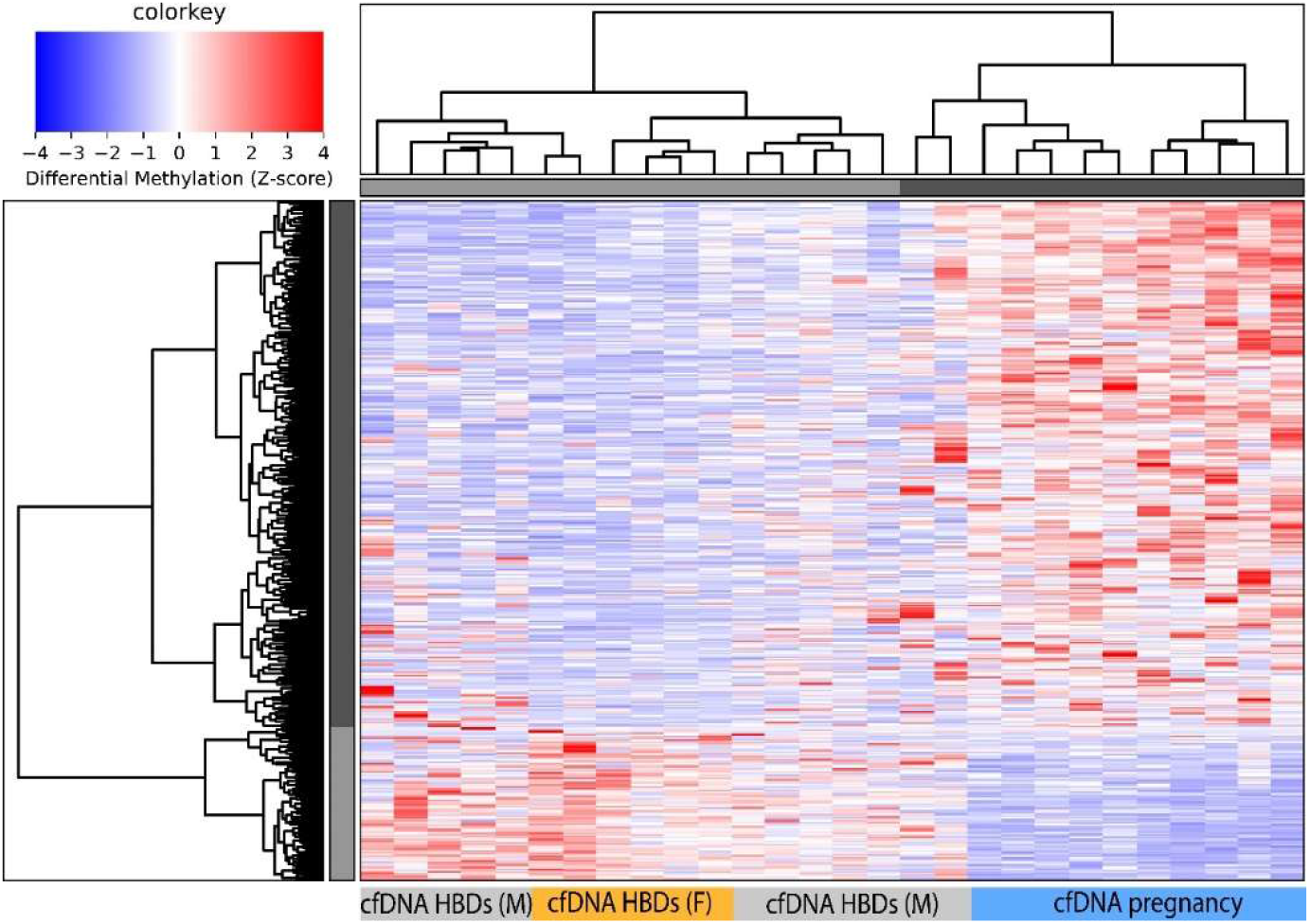
Heatmap visualizing overlap between cfDNA from male (M) and female (F) non-pregnant healthy blood donors (HBDs) compared to cfDNA from pregnant women. Selected DMRs identified between pregnant women (P) and non-pregnant female HBDs. Red represents hypermethylation, blue represents hypomethylation.

**Supplemental Figure2.**
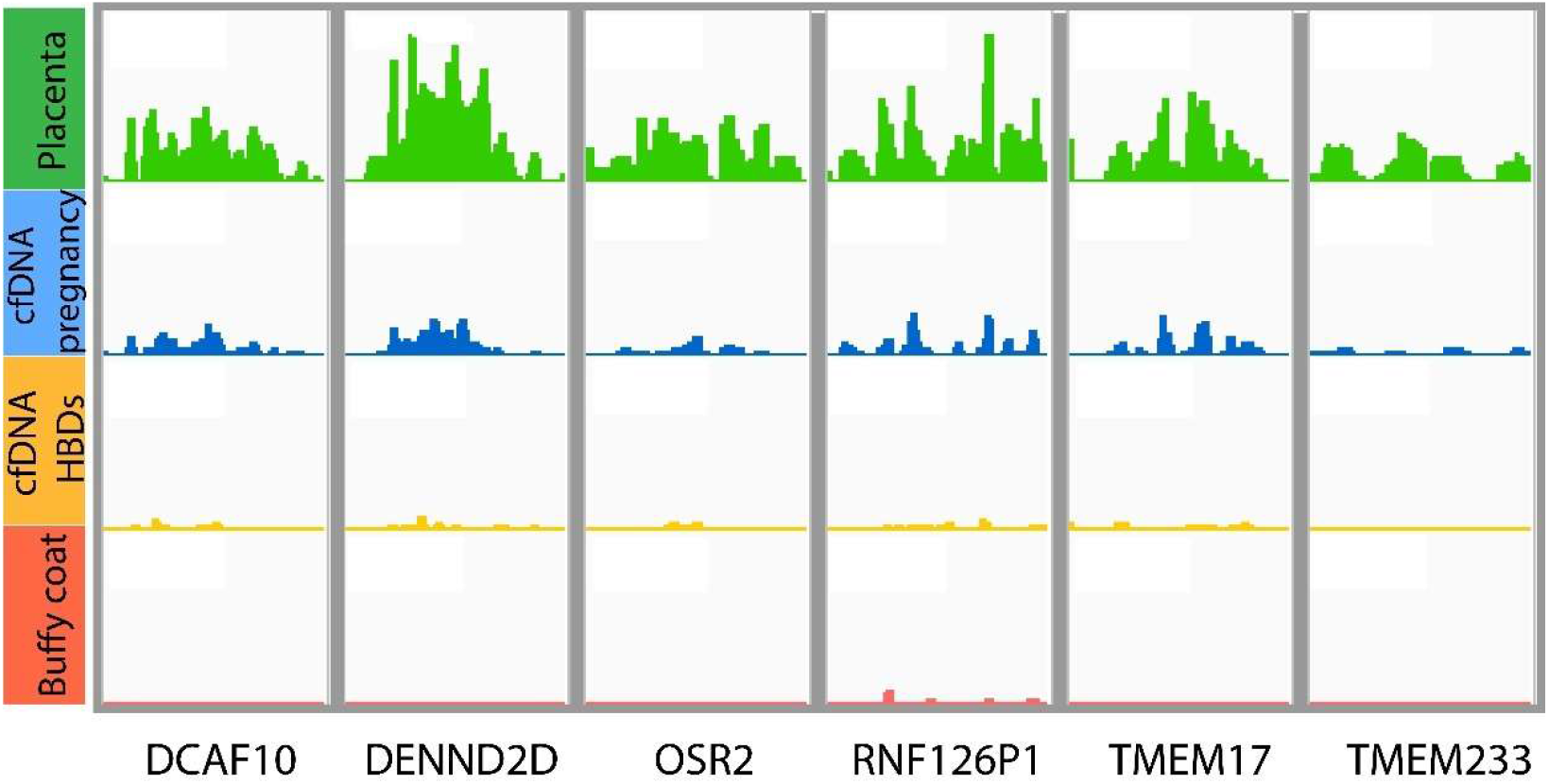
Gene-tracks for DMRs between cfDNA from pregnant women and non-pregnant female healthy blood donors (HBDs) that overlap with CpGs previously identified by Chu et al (30).

**Table S1**: Overview of used samples and their technical performance.

**Table S2**: DMRs identified between cfDNA from pregnant women compared to non-pregnant women

**Table S3**: Cumulative methylation scores for all included samples for DMRs identified in cfDNA between pregnant and non-pregnant women.

**Table S4**: *RASSF1* promoter methylation in placental tissue biopsies and in cfDNA from pregnant and non-pregnant women.

**Table S5**: Identified Y-chromosomal regions in cfDNA from male compared to female HBDs. Identified DMRs are applied in cfDNA from pregnant women to determine fetal sex.

**Table S6**: Baseline characteristics of women with buffy coat samples (n=10). Categorical variables are described using absolute numbers and percentages (%). Continuous variables are described using medians and interquartile ranges (IQR). IVF/ICSI = in vitro fertilization/intracytoplasmic sperm injection.

